# The development of individual differences in cooperative behaviour: maternal glucocorticoid hormones alter helping behaviour of offspring in wild meerkats

**DOI:** 10.1101/316182

**Authors:** Ben Dantzer, Constance Dubuc, Ines Braga Goncalves, Dominic L. Cram, Nigel C. Bennett, Andre Ganswindt, Michael Heistermann, Chris Duncan, David Gaynor, Tim H. Clutton-Brock

## Abstract

The phenotype of parents can have long-lasting effects on the development of offspring as well as on their behaviour, physiology, and morphology as adults. In some cases, these changes may increase offspring fitness but, in others, they can elevate parental fitness at a cost to the fitness of their offspring. We show that in Kalahari meerkats (*Suricata suricatta*), the circulating glucocorticoid (GC) hormones of pregnant females affect the growth and cooperative behaviour of their offspring. We performed a 3-year experiment in wild meerkats to test the hypothesis that GC-mediated maternal effects reduce the potential for offspring to reproduce directly and therefore cause them to exhibit more cooperative behaviour. Daughters (but not sons) born to mothers treated with cortisol during pregnancy grew more slowly early in life and exhibited significantly more of two types of cooperative behaviour (pup rearing and feeding) once they were adults compared to offspring from control mothers. They also had lower measures of GCs as they aged, which could explain the observed increases in cooperative behaviour. Because early life growth is a crucial determinant of fitness in female meerkats, our results indicate that GC-mediated maternal effects may reduce the fitness of offspring, but may elevate parental fitness as a consequence of increasing the cooperative behaviour of their daughters.

## Introduction

Parental effects are a mechanism of trans-generational phenotypic plasticity that occurs when the parental phenotype or parental environment modifies offspring characteristics (1). Parental effects can increase the survival or reproduction of offspring, thereby elevating the direct fitness of both offspring and parents (2-6). Alternatively, parental effects can increase parental fitness, but at some cost to the fitness of their offspring (7-8) – a process regarded as a type of parental manipulation (9-12) or ‘selfish parental effect’ (13). For example, in mammals, the optimal birth weight or litter size often differs between mothers and offspring (14) and pregnant females experiencing stressful environments may reallocate resources away from offspring and towards themselves, so that their offspring are smaller or grow more slowly before weaning (15). Despite these observations, it has been suggested that selfish parental effects may be rare and unstable because selection would be expected to favour the evolution of resistance mechanisms in offspring (7, 11, 13, 16, 17).

Selfish parental effects may in fact be more likely in cooperatively breeding species where philopatric offspring (subordinates) help to rear the subsequent offspring of their parents or other close relatives. This could be especially likely under low food or high stress conditions as parents may gain substantial direct fitness benefits from delaying the development of their offspring if this causes them to invest in alloparental care directed at the parent’s subsequent offspring (9-10). In addition, the costs of selfish parental effects to offspring could be reduced in these circumstances, as offspring will gain indirect fitness benefits by contributing to raising the subsequent offspring of their parents (18). For example, laboratory studies of eusocial insects suggest the possibility that selection will favour the evolution of alleles that enable mothers to increase the helping behaviour of their offspring while simultaneously reducing their capabilities of reproducing on their own (19-20; but see 21).

To date, empirical field tests of how parental effects shape the helping behaviour of offspring are rare (23) and studies of selfish parental effects have mostly focused on non-social species (13, 24). Here, we report the results of experiments designed to test the hypothesis that elevated maternal glucocorticoid levels (GCs) reduce the potential for offspring to have direct reproductive opportunities and causes them to exhibit more cooperative behaviour. In a 3-year field study, we experimentally elevated maternal GCs by treating pregnant dominant female meerkats with cortisol and tracking the growth, stress physiology, and cooperative behaviour of their offspring from birth until ~18 months of age, compared to those from control litters. We manipulated maternal GCs because they are known to cause mothers to reallocate energy away from offspring and towards themselves (15), indicating that they may function as a mediator of selfish maternal effects. Changes in maternal GCs have also previously been shown to delay the dispersal of offspring as well as influence the parental care behaviour of offspring (25-26), both traits that are important in cooperative breeders where philopatric offspring exhibit alloparental care behaviour towards juveniles.

To identify if the exposure of mothers to heightened GCs reduced reproductive success of their offspring, we examined if offspring from mothers treated with cortisol during pregnancy grew more slowly early in life. In meerkats, the rate of early life growth and body mass is closely linked to future direct fitness through its effects on survival, foraging success, adult body mass (27-29), as well as the probability of acquiring dominance (30-31) and other direct reproductive opportunities (32). As elevated exposure to maternal GCs in some mammals may reduce offspring size and growth early in life (15), we predicted that offspring from mothers treated with cortisol during pregnancy would be smaller or grow more slowly early in life. Because the rate of early life growth is predictive of future direct fitness in meerkats (27-32), we predicted that if offspring from mothers treated with cortisol did grow more slowly, they would consequently invest more in indirect fitness opportunities by contributing more to cooperative activities than controls.

Secondly, we determined if offspring from mothers treated with cortisol during pregnancy subsequently increased their contributions to two types of cooperative behaviours: pup rearing (“babysitting”: 33) and food provisioning during the period when the pups are foraging with their natal group, but are not yet nutritionally independent (“pup feeding”: 34). We chose these two behaviours as they appear to be most costly from an energetic perspective (35) and are most closely tied to the probability of parents successfully rearing offspring. If offspring from mothers treated with cortisol during pregnancy exhibit more of either of these two types of alloparental care, this should increase both parental direct fitness (the number of offspring that they subsequently produce) and the indirect fitness of offspring, because subsequent offspring that receive more alloparental care should grow faster or have higher early life survival (27, 32, 36, 37). Previous studies in meerkats show that offspring with more helpers or those that receive more alloparental care grow faster or have early life survival (27, 32, 36, 37).

To assess the mechanism by which elevated exposure to maternal stress may affect the alloparental care behaviour of offspring, we repeatedly measured plasma cortisol and faecal glucocorticoid metabolite (fGCM) concentrations of offspring from when they were approximately 1 to 18 months of age to identify how our manipulations affected their neuroendocrine stress axes (GC output). Elevated maternal GCs can cause long-term changes in the neuroendocrine stress axis of offspring (38) and elevated activity of the neuroendocrine stress axis in meerkats can reduce their contributions to alloparental care (39). We therefore predicted that if offspring born to mothers treated with cortisol during pregnancy exhibited more alloparental care behaviour compared to controls, they would also have reduced plasma cortisol and fGCM concentrations.

## Methods

### Study site & basic data collection

We studied free-living meerkats at the Kuruman River Reserve (26° 58’ S, 21° 49’ E) in the Northern Cape, South Africa from 2014-2017. Individuals were marked uniquely with PIT tags (Identipet^®^, Johannesburg, South Africa) as well as dye marks so that they could be identified. Groups were visited for ~4-8 hours per day ~4-6 times per week throughout each year of study and sometimes more frequently such as when there were pups being babysat. Groups were visited at sunrise before meerkats emerged from their sleeping burrow. After all the meerkats had emerged, but prior to when they started going foraging, we counted the total number of meerkats in the group (to get estimates of group size) and recorded which individuals were present (using their unique combinations of dye marks). We recorded their body mass on a portable balance each morning before foraging, 2-4 hrs after foraging was initiated, and immediately prior to when foraging ended (40). These measures of body mass provided our estimates of growth, body mass, and foraging success that are used in our analyses described below.

### Experimental manipulations of dominant females

Dominant females in each group were identified via behavioural observations (41). The pregnancy status of dominant females was determined visually (distended abdomen) as well as noting a constant increase in their body mass. Dominant females were treated with either a cortisol solution or a control oil vehicle when they were pregnant by feeding them food containing one of these two treatments. We initially offered experimental animals hard boiled eggs with added cortisol but found that they rejected all foods that contained added cortisol with the exception of scorpions. We consequently fed experimental females with cortisol (10 mg/kg of hydrocortisone, Sigma H4126), that were dissolved in 100 μl of 100% coconut oil and injected into a dead scorpion (*Opistophthalmus* spp.). Control females were fed a dead scorpion that was injected with 100 μl of 100% coconut oil. A previous study using the same protocol showed that meerkats that were fed cortisol had significantly higher plasma cortisol and fGCM concentrations than control animals and these increases were within a biologically relevant range (39). This indicated that our treatment causes the exogenous glucocorticoids that we feed the meerkats to enter their bloodstream and leads to sustained increases in their circulating glucocorticoid concentrations.

Females were randomly allocated to the treatments. Across the three years of this study, we produced a total of 13 cortisol-treated litters from 10 females and 7 control litters produced by 6 females (Table 1). Three of the females experienced both the control and cortisol treatments at different time points of the experiment, whereas one female experienced the control treatment once and the cortisol treatments twice. For these latter females treated twice, the order of treatments was randomly selected. We conducted these experiments over the course of three years: 13 litters from 10 females in 2014 (April-December 2014, 3 litters aborted), 5 litters from 5 females in 2015 (February-July 2015, 1 litter aborted), and 3 litters from 3 females in 2016 (July 2016). Three cortisol-treated mothers and one control mother aborted their offspring prior to birth and were excluded from any analyses except for assessing differences in the frequency of abortion between control and cortisol-treated mothers (Table 2). This provided final sample sizes of 31 pups from 10 litters from 9 cortisol-treated females and 25 pups from 6 litters from 6 control females (Table 1).

**Table 1.** Summary of effects of dominant female treatments on litter characteristics and offspring survival. Number of pups emerged correspond to those that emerged from the natal burrow and in some cases these pups died before their sex could be determined (shown as “Unk”). Three of the 13 litters treated with cortisol and one of the 7 control (fed) litters were aborted prior to birth.

| Treatment | Total # litters<br>& females<br>treated | Total # pups<br>emerged<br>(F, M, Unk) | Avg. Pups<br>emerged | Avg. Pups<br>Surviving to<br>3 months | Avg. Pups<br>Surviving to<br>6 months | Avg. Pups<br>Surviving to<br>12 months |
| --- | --- | --- | --- | --- | --- | --- |
| Untreated | 52 (21 females) | 49 F, 84 M, 52 Unk | $3.78 \pm 1.23$ | $2.68 \pm 1.58$ | $2.1 \pm 1.63$ | $1.62 \pm 1.54$ |
| Control | 7 (6 females) | 12 F, 13 M | $4.17 \pm 0.98$ | $3.83 \pm 1.17$ | $3.83 \pm 1.17$ | $2.83 \pm 1.94$ |
| Cortisol | 13 (10 females) | 13 F, 18 M | $3.87 \pm 0.83$ | $3.25 \pm 1.03$ | $2.75 \pm 1.28$ | $1.75 \pm 1.75$ |

**Table 2.** Effects of dominant female treatments (cortisol or control) on litter characteristics and pup survival. Results are from a linear mixed-effects model (# pups emerged) or generalized linear mixed-effects models (GLMMs, all other response variables) that each contained random intercept terms for dominant female identity and year. No GLMM was overdispersed as indicated by goodness of fit tests (R package aods3, P-values from Pearson χ^2^ tests ranged from 0.13 to 1). The number of litters aborted by untreated females was not known so we only assessed the effects of cortisol vs. control treatments on the number of litters aborted. Litter sex ratio is the proportion of males in the litter.

| Response variable | Fixed effect | b | SE | t or z | P-value |
| --- | --- | --- | --- | --- | --- |
| **# Litters aborted** |  |  |  |  |  |
|  | Intercept | -1.21 | 0.66 | -1.83 | 0.07 |
|  | Birthdate | -0.17 | 0.59 | -0.29 | 0.77 |
|  | Treatment |  |  |  |  |
|  | *Control* | -0.59 | 1.27 | -0.46 | 0.64 |
| **# Pups emerged** |  |  |  |  |  |
|  | **Intercept** | **3.9** | **0.42** | **9.36** | **<0.0001** |
|  | Birthdate | 0.02 | 0.16 | 0.11 | 0.91 |
|  | Treatment |  |  |  |  |
|  | *Control* | 0.3 | 0.62 | 0.49 | 0.63 |
|  | *Untreated* | -0.11 | 0.45 | -0.25 | 0.8 |
| **Litter sex ratio** |  |  |  |  |  |
|  | Intercept | -0.54 | 0.3 | -1.84 | 0.066 |
|  | Birthdate | 0.02 | 0.12 | 0.15 | 0.88 |
|  | Treatment |  |  |  |  |
|  | *Control* | -0.11 | 0.45 | -0.23 | 0.81 |
|  | *Untreated* | 0.09 | 0.33 | 0.27 | 0.78 |
| **Prop. litter surviving to 3 months** |  |  |  |  |  |
|  | Intercept | -0.16 | 0.27 | -0.62 | 0.53 |
|  | Birthdate | -0.12 | 0.11 | -1.09 | 0.27 |
|  | Treatment |  |  |  |  |
|  | *Control* | 0.06 | 0.39 | 0.16 | 0.87 |
|  | *Untreated* | -0.19 | 0.3 | -0.64 | 0.52 |
| **Prop. litter surviving to 6 months** |  |  |  |  |  |
|  | Intercept | -0.32 | 0.28 | -1.16 | 0.24 |
|  | Birthdate | -0.19 | 0.11 | -1.65 | 0.099 |
|  | Treatment |  |  |  |  |
|  | *Control* | 0.21 | 0.4 | 0.52 | 0.6 |
|  | *Untreated* | -0.28 | 0.31 | -0.91 | 0.36 |
| **Prop. litter surviving to 12 months** |  |  |  |  |  |
|  | **Intercept** | **-0.91** | **0.46** | **-1.999** | **0.046** |
|  | Birthdate | -0.22 | 0.14 | -1.58 | 0.11 |
|  | Treatment |  |  |  |  |
|  | *Control* | 0.36 | 0.46 | 0.79 | 0.43 |
|  | *Untreated* | 0.096 | 0.38 | 0.25 | 0.8 |

Reference for Treatment was cortisol-treated mothers. Data other than # litters aborted are based upon an initial sample size of offspring from untreated (195 pups from 52 litters produced by 21 dominant females), control (25 pups from 6 litters produced by 6 females), or cortisol-treated (31 pups from 10 litters produced by 9 females) litters that produced pups that emerged from the burrow.

We aimed to experimentally increase the glucocorticoid concentrations of pregnant dominant females from when they were first confirmed to be pregnant (second half of gestation) until parturition. Gestation in meerkats is ~70 days so we aimed to treat them with glucocorticoids from approximately 35-70 d during gestation. In reality, females that successfully produced a litter where pups emerged from the natal burrow were treated with cortisol for 12-36 days prior to birth (n=10 litters from 9 females, mean = 23.7 d, median = 23.5 d), whereas controls were fed for 12-58 d prior to birth (n=6 litters from 6 females, mean = 30 d, median = 20.5 d). Although controls were treated for slightly longer, there was no significant difference in treatment duration between control and cortisol-treated females (general linear model, t = 1.05, P = 0.31).

To provide an additional comparison group to investigate how our treatments (fed during pregnancy or fed cortisol during pregnancy) affected offspring survival, growth, and cooperative behaviour, we also monitored these traits in offspring produced by dominant females that were untreated during pregnancy (n = 52 litters from 21 dominant females, Table 1). These females were not fed or treated with cortisol (hereafter, “untreated mothers”). For our analyses of how the treatments affected offspring survival and growth, the untreated offspring were those from litters produced by dominant females in other meerkat groups in our same study area and were born during our study. We assessed the contributions of offspring from mothers treated with cortisol during pregnancy to two cooperative behaviours (babysitting and pup feeding) compared to those from control mothers, but also to other group members from untreated mothers. We did not have data from offspring from untreated mothers when we assessed how our treatments affected their plasma cortisol or fGCM concentrations.

### Quantifying early life growth of offspring

Meerkat pups typically first emerge from their natal burrow approximately 21-30 d after birth. Meerkat groups and dominant females were monitored daily around the estimated date of parturition and birth dates were estimated according to the change in the physical appearance of the dominant female, a large drop in body mass overnight, and group members exhibiting babysitting behaviour at the sleeping burrow. Burrows containing pups were monitored each day and, when pups emerged, they were uniquely marked by trimming small sections of hair before permanent PIT tags could be applied. Pups were weighed each time we visited the groups on a portable balance in the morning after group members emerged from their sleeping burrow (as above).

### Quantifying cooperative behaviour of offspring

We measured the babysitting (controls: 195-655 d; cortisol: 184-655 d; untreated: 155-655 d) and pup feeding (controls: 220-635 d; cortisol: 184-655 d; untreated: 155-626 d) contributions of offspring from cortisol-treated and control mothers when they were >6 months of age until death or disappearance. We visited sleeping burrows containing pups every day in the morning and recorded the identity of the attending babysitters. As we have done previously (33, 36, 39, 42), we calculated relative babysitting contributions of each individual meerkat for each litter by dividing the total number of days an individual babysat a litter over the total number of days that this specific litter had a babysitter. Pup feeding behaviour for each pup produced by the dominant females in the different treatment groups was estimated using *ad libitum* sampling (34, 39). When the social group contained pups (up to 90 d of age), we recorded all pup-feeding events from all individuals, which are visually and acoustically conspicuous to observers (43). We then used these data to estimate the proportion of pup-feeding events exhibited by an individual relative to all others in the group (i.e., relative pup feeding). Because the total amount of time devoted to the *ad libitum* recording sessions varied, we corrected for variation in observation time (see below).

### Quantifying plasma cortisol concentrations from offspring

We obtained plasma samples from offspring from cortisol-treated and control mothers approximately every 3 months from first emergence from the burrow (~1 month) until ~18 months of age (controls: 20-548 d; cortisol-treated: 25-559 d). Capture and blood processing procedures are described elsewhere (44-45). The amount of time it took to obtain the blood samples varied (median = 10.6 min, SD = 7.2 min), but we included co-variates for sampling time and sampling time^2^ to control for effects of sampling time (described in 45). We measured total plasma cortisol concentrations using a previously validated assay (Coat-a-Count, Siemens Diagnostic Products Corporation, Los Angeles, USA: validation described in 44). The sensitivity of the assay was 1.9 ng/ml and cross-reactivity to other hormones was 76% with prednisolone, 11.4% with 11-deoxycortisol, 2.3% with prednisone and <1% with aldosterone, corticosterone, cortisone, oestriol, estrone and pregnenolone. Intra-assay coefficient of variation (CV) was 7% (n = 20 samples). Inter-assay CV for a low control (78.5 ± 6.3 ng/ml n = 5 assays) was 8% and 2.8% for a high control (187 ± 5.3 ng/ml, n = 5 assays).

### Quantifying fGCM concentrations from offspring

We collected faecal samples from offspring of cortisol-treated and control mothers opportunistically during behavioural observations over the course of the study (controls: 25-356 d; cortisol-treated: 32-326 d). Faecal samples were processed as described previously using a methanol solution to extract fGCMs for analysis (46-47). Immunoreactive fGCM concentrations were determined using a group-specific enzyme immunoassay measuring cortisol metabolites with a 5β-3α,11β-diol-structure (11β-hydroxyetiocholanolone), already validated and established for monitoring fGCM alterations in meerkats (47). Faecal GCMs measured reflect average adrenal cortisol production over the previous ~24 to 48 hr period (47). Detailed assay characteristics, including full descriptions of the assay components and cross-reactivities, are found elsewhere (48). The sensitivity of the assay was 1.2 ng/g dry weight and intra-assay CV determined by repeated measurements of high and low value quality controls were 6.9% and 7.4% and inter-assay CV values were 11.5% and 15.9% (n = 29 assays), respectively.

### Statistical analyses

We used generalized (binomial errors) or linear mixed-effects models (LMMs) to examine how our treatments affected the probability that the litter was aborted, litter size and sex ratio at emergence from the burrow, and the proportion of the litter that survived to emergence from the burrow, independence (~90 d of age: 29), and 6 or 12 months of age. We focused on addressing whether the offspring from cortisol-treated mothers differed from control or untreated mothers. These models included a fixed effect for date of birth of the litter and random intercept terms for dominant female identity and year (as the experiments were conducted over 3 years). None of the GLMMs were overdispersed (Table 2).

We used a LMM to investigate how the maternal treatments affected offspring growth from first emergence from their natal burrow (~1 month) to 3 months of age when the pups are typically foraging independently (29, 34). Morning body mass (in grams) was the response variable with the fixed effects of maternal treatment (cortisol-treated, control, or untreated), pup sex, pup age, litter size at burrow emergence, first measure of body mass when the pups first emerged from the burrow (to control for possible differences in age or development when they entered our study population), group size, group size^2^, total rainfall in the previous 60 days, two measures of seasonality (sine and co-sine functions of day of weight measure: see 40), and two three-way interactions between sex, treatment, age or age^2^. Group size was defined as the average number of subordinate meerkats >6 months of age in the group during the entire period of offspring growth. Random intercept terms for year and the identity of the individual nested in litter, nested in dominant female identity, nested in group were also included in this model. Fixed and random effects included in these models were based upon previous studies investigating meerkat body mass and/or growth from 1-3 months (28, 31, 40). To prevent any issues associated with selective disappearance of specific individuals, only individuals that survived to 90 d were included in these analyses.

We assessed how the treatments affected the relative babysitting and pup feeding contributions of subordinates when they were >6 months (as they rarely do alloparental care behaviour when <6 months: 36) from cortisol-treated, control, and untreated mothers. Relative babysitting and pup feeding contributions are defined as the proportion of babysitting or pup feeding contributions exhibited by a specific individual compared to the total number of babysitting or pup feeding contributions for that litter exhibited by all individuals in the group that were >6 months of age at the time of the birth of the litter (36, 39, 42). In these generalized linear mixed-effects models (GLMM, binomial errors), we included a three-way interaction between treatment, sex, and age of the subordinate to assess if the effects of the treatments on babysitting or pup feeding varied according to the sex or age of the subordinate, as contributions to cooperative behaviour in meerkats are known to vary according to subordinate sex and age (36). To account for differences in observation time, we included a co-variate for the number of days the litter was babysat (babysitting length) and the number of days the subordinate was observed in the group during babysitting as well as the total time spent observing the group during pup feeding (observation time). We included a range of co-variates (see Tables 4-6) that have been previously documented to affect relative contributions to babysitting and pup feeding, including age, foraging success, body mass, and group size (34-36, 42, 49; 50). Group size was defined as the average number of subordinate meerkats >6 months of age in the group while the litter was being babysat or pup fed. Foraging success was defined as the average weight gained per hour estimated as the change in body mass from morning weight to evening weight over the total number of hours that had elapsed since those two weights (45). Relatedness between the subordinate and the litter being babysat was not included as it has not been shown to impact babysitting or pup feeding contributions (27, 42) and nearly all of the litters in our dataset were produced by the mother or full sibling of the subordinate. Random intercept terms for year and the identity of the individual, and litter being babysat or pup fed were nested within the group where the litter was being babysat or pup fed. Overdispersion was not an issue for our GLMM for babysitting as indicated by the goodness of fit test (Pearson χ^2^ = 147.1, df = 154, P = 0.64, using package aods3: 51) but our GLMM for pup feeding was initially overdispersed (Pearson χ^2^ = 310, df=165, P < 0.0001) so we included an observation level random intercept term.

We used two separate LMMs to assess how our manipulations affected plasma cortisol and fGCMs in offspring from cortisol-treated and control mothers (we did not have these data from offspring from untreated mothers). Each model included fixed effects for maternal treatment, pup sex and age, time of day and year that the sample was acquired (2014 or 2015), and random intercept terms for identity of individual nested in their birth litter and group. In the model for plasma cortisol concentrations, we also included a linear and second order fixed effect for the time it took to acquire the blood sample to control for any variation in plasma cortisol concentrations due to restraint stress (45). Year was included as a fixed effect because we only had samples from two separate years. We included covariates associated with the individual meerkat and weather or social group characteristics that are known to affect plasma cortisol (45) or fGCM (47) concentrations (see Tables 6-7).

We used R (version 3.4.3: 52) for all of our statistical analyses. R package lme4 (version 1.1-14: 53) was used for LMMs and P values were estimated using lmerTest (version 2.0-33: 54). A graphical approach was used to confirm normality and homoscedasticity of residuals and to confirm there were no observations with high leverage (55). Collinearity among predictor variables included in our models was assessed by calculating variance inflation factors (55) or generalized variance inflation factors (for variables that had a second order term or those included in an interaction: 56). Collinearity was not a problem as indicated by our variance inflation factors (VIFs) as VIFs or generalized VIFs were less than ~4 for all variables. In our model for how our treatments affected offspring growth (Table 3), the generalized VIF for the two measures of seasonality (sine and co-sine functions of day of weight measure) were <6 but these two variables were included a priori given their previously documented effects on body mass and growth in meerkats (40). All continuous variables were standardized to a mean of 0 and SD of 1.

**Table 3.** Effect of dominant female treatments on offspring growth from emergence to nutritional independence (1-3 months of age). Data are from a linear mixed-effects model where the response variable was morning body mass that contained random intercept terms for individual identity nested in birth litter nested in mother nested in natal group (σ^2^ = 116.7) and year (σ^2^ = 0). If fixed effects by themselves were involved in significant higher order interactions with other variables, only parameter estimates are shown.

| Fixed Effect | b | SE | t | df | P-value |
| --- | --- | --- | --- | --- | --- |
| **Intercept** |  |  |  |  |  |
| *Females* | **184.6** | **9.75** | **18.93** | **53** | **<0.0001** |
| *Males* | **189.1** | **9.5** | **19.8** | **49** | **<0.0001** |
| Litter size | -5.31 | 3.9 | -1.37 | 48 | 0.18 |
| **First weight** | **15.96** | **2.52** | **6.344** | **166** | **<0.0001** |
| Sex (M) | 4.54 | 4.32 |  |  |  |
| Age | 53.3 | 1.06 |  |  |  |
| Age<sup>2</sup> | -4.76 | 1.15 |  |  |  |
| **Rainfall** | **2.81** | **0.61** | **4.57** | **4550** | **<0.0001** |
| **Season (Sine)** | **-67.2** | **3.3** | **-20.6** | **4238** | **<0.0001** |
| **Season (Co-sine)** | **66.22** | **3.4** | **19.2** | **4191** | **<0.0001** |
| Group size | 2.3 | 2.66 | 0.86 | 109 | 0.39 |
| Group size<sup>2</sup> | -0.09 | 2.21 | -0.04 | 178 | 0.97 |
| Treatment (Control) |  |  |  |  |  |
| *Females* | 22.6 | 15.6 |  |  |  |
| *Males* | 12.9 | 15.5 |  |  |  |
| Treatment (None) |  |  |  |  |  |
| *Females* | 23.6 | 10.6 |  |  |  |
| *Males* | 22.9 | 10.4 |  |  |  |
| Age x Sex (M) | 1.64 | 1.37 | 1.19 | 4567 | 0.23 |
| Age<sup>2</sup> x Sex (M) | 1.02 | 1.53 | 0.67 | 4543 | 0.5 |
| Sex (M) x Treatment (Control) | -9.73 | 7.7 | -1.27 | 164 | 0.21 |
| Sex (M) x Treatment (None) | -0.63 | 5.1 | -0.12 | 172 | 0.9 |
| **Age x Treatment (Control)** |  |  |  |  |  |
| *Females* | **7.6** | **1.8** | **4.17** | **4595** | **<0.0001** |
| *Males* | -2.3 | 1.5 | -1.48 | 4548 | 0.14 |
| Age x Treatment (None) |  |  |  |  |  |
| *Females* | 0.79 | 1.2 | 0.65 | 4574 | 0.51 |
| *Males* | -0.53 | 1.01 | -0.52 | 4580 | 0.6 |
| Age<sup>2</sup> x Treatment (Control) |  |  |  |  |  |
| *Females* | -0.13 | 1.93 | -0.07 | 4521 | 0.94 |
| *Males* | -1.05 | 1.6 | -0.65 | 4547 | 0.52 |
| **Age<sup>2</sup> x Treatment (None)** |  |  |  |  |  |
| *Females* | -2.6 | 1.3 | -1.96 | 4534 | 0.051 |
| *Males* | **-6.9** | **1.1** | **-6.23** | **4556** | **<0.0001** |
| **Age x Treatment (Control) x Sex (M)** | **-9.9** | **2.3** | **-4.23** | **4574** | **<0.0001** |
| Age x Treatment (None) x Sex (M) | -1.3 | 1.6 | -0.84 | 4581 | 0.4 |
| Age<sup>2</sup> x Treatment (Control) x Sex (M) | -0.93 | 2.5 | -0.37 | 4552 | 0.71 |
| **Age<sup>2</sup> x Treatment (None) x Sex (M)** | **-4.3** | **1.7** | **-2.5** | **4552** | **0.01** |

Data used in these analyses were 4676 measures of body mass from 195 meerkats produced by 21 dominant females across 53 different litters in 16 different social groups in three different years. Only offspring that survived to 90 days of age were included in these analyses.

## Results

### Effects of treatments on litter characteristics and offspring survival

There was no evidence that the treatment of pregnant females with cortisol affected their ability to maintain litters to term or the survival of their pups prior to emergence from the natal burrow (Tables 1-2). The number of pups surviving to emergence from the natal burrow or 3, 6, or 12 months of age and the litter sex ratio were not different among litters from cortisol-treated, control, or untreated females (Tables 1-2).

### Effects of treatments on offspring early life growth

The effects of the treatments on offspring growth from initial emergence to nutritional independence (1-3 months) differed between daughters and sons, as reflected in the significant three-way interaction between treatment, sex, and age (Table 3). Daughters (but not sons) from cortisol treated mothers grew more slowly from 1-3 months compared to those from control (fed) mothers (daughters: age x treatment, *t* = - 4.17, *P* < 0.0001; sons: *t* = −1.48, *P* = 0.14), but exhibited similar growth compared to those from untreated (unfed) mothers (daughters: age x treatment, *t* = 0.65, *P* = 0.51; sons: *t* = −0.52, *P* = 0.6, Table 3, Fig. 1). Daughters, but not sons, from control (fed) mothers grew faster than those from untreated mothers (daughters: age x treatment, *t* = −4.24, P < 0.0001; sons: *t* = 1.35, *P* = 0.18).

**Figure 1.**
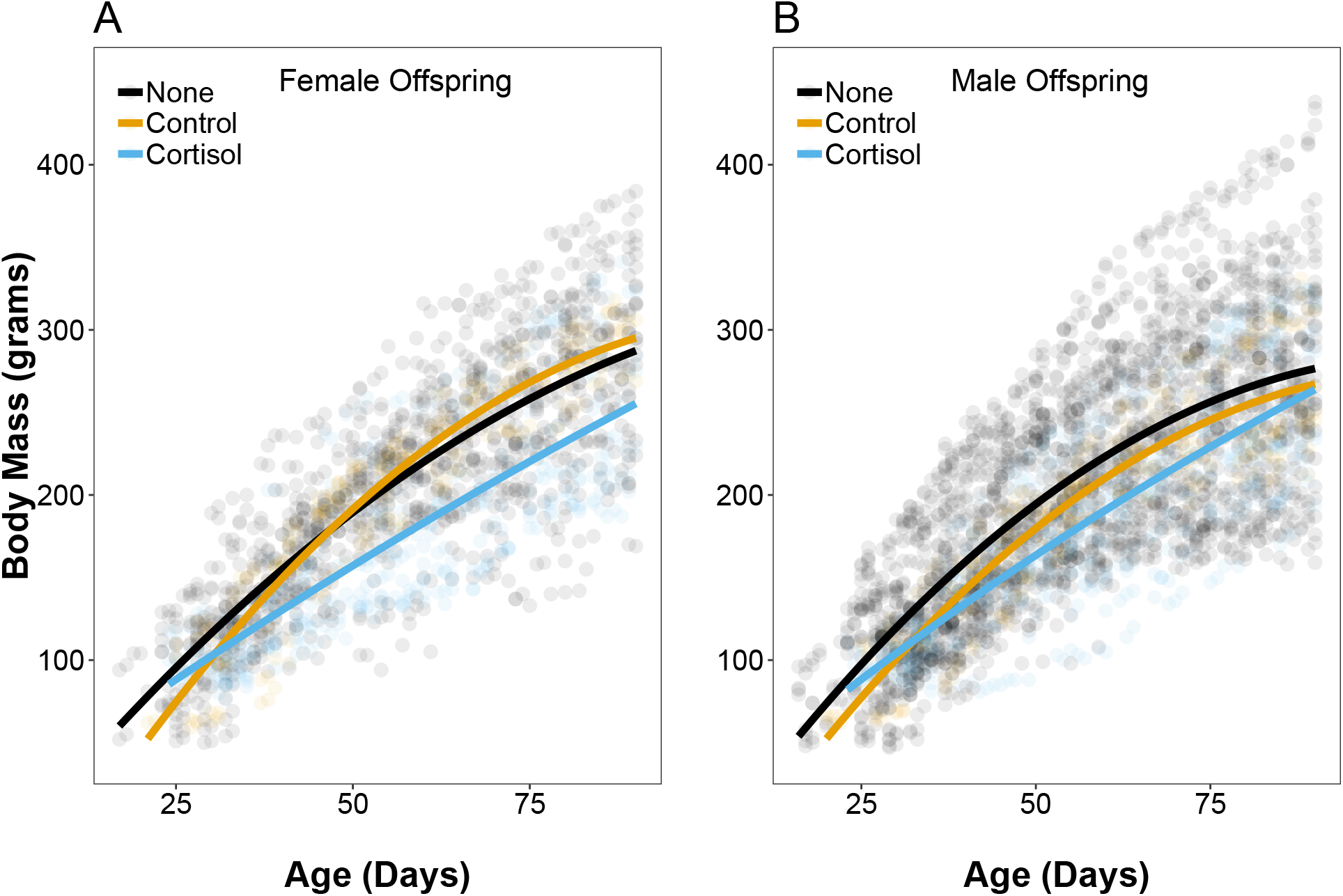
(A) Daughters but not (B) sons from mothers treated with cortisol during pregnancy were significantly smaller from initial emergence from their natal burrow to nutritional independence (~1-3 months) compared to those from control mothers (daughters: age x treatment, *t* = −4.17, *P* < 0.0001; sons: *t* = −1.48, *P* = 0.14), but not untreated mothers (daughters: age x treatment, *t* = 0.65, *P* = 0.51; sons: *t* = −0.52, *P* = 0.6, Table 3). Data are body mass measures from offspring from cortisol-treated (females: n = 373 estimates; males: n = 488), control (females: n = 215; males: n = 241), and untreated mothers (females: n = 1121; males: n = 2238). Raw data and regression lines are shown (full results in Table 3).

### Effects of treatments on offspring cooperative behaviour

The effects of the maternal treatments on babysitting behaviour of offspring depended upon the age and sex of the offspring (Table 4). Babysitting contributions in daughters from mothers treated with cortisol during pregnancy were slightly, but significantly higher with increasing age of the babysitter compared to those from control mothers (age x treatment, *z* = −2.89, *P* = 0.0039) but not untreated mothers (age x treatment, *z* = 1.88, *P* = 0.06; Table 4, Fig. 2). Babysitting contributions in sons from mothers treated with cortisol during pregnancy showed a similar tendency to slightly increase with age compared to those from control mothers, but this difference was not significant (age x treatment, *z* = −1.92, *P* = 0.055). Further, age-related increases in babysitting contributions between males from mothers treated with cortisol during pregnancy and untreated mothers did not differ (age x treatment, *z* = −0.03, *P* = 0.97; Table 4, Fig. 2). Comparisons of the magnitude of effect sizes showed that the interaction between age and maternal treatment had a larger effect on babysitting contributions in daughters but not sons than other variables known to impact babysitting contributions, such as foraging success, age-related body mass, or group size (Table 4).

**Table 4.** Effect of dominant female treatments on relative babysitting contributions. Data are from a generalized linear mixed-effects model where the response variable is the proportion of babysitting exhibited by the subordinate meerkat relative to the total babysitting contributions the litter received. The model contained random intercept terms for individual (σ^2^ = 0.12), litter nested within group (σ^2^ = < 0.0001), and year (σ^2^ = 0.000). If fixed effects by themselves were involved in significant higher order interactions with other variables, only parameter estimates are shown.

| Fixed Effect | b | SE | z | P-value |
| --- | --- | --- | --- | --- |
| **Intercept** |  |  |  |  |
| *Females* | **-2.1** | **0.22** | **-9.37** | **< 0.0001** |
| *Males* | **-2.14** | **0.19** | **-11.01** | **< 0.0001** |
| **Babysitting length** | **0.24** | **0.08** | **2.77** | **0.0056** |
| **Observation time** | **-0.28** | **0.08** | **-3.4** | **0.0007** |
| Litter size | 0.015 | 0.05 | 0.32 | 0.75 |
| Mixed Litter? | -0.04 | 0.12 | -0.31 | 0.76 |
| Sex (M) | -0.03 | 0.27 | -0.13 | 0.9 |
| Age |  |  |  |  |
| *Females* | 0.13 | 0.17 |  |  |
| *Males* | 0.4 | 0.19 |  |  |
| Foraging success | -0.05 | 0.05 | -0.99 | 0.32 |
| Mass |  |  |  |  |
| *Females* | -0.32 | 0.12 |  |  |
| *Males* | -0.098 | 0.14 |  |  |
| **Group size** |  |  |  |  |
| *Females* | **-0.35** | **0.09** | **-3.97** | **< 0.0001** |
| *Males* | **-0.27** | **0.08** | **-3.28** | **0.001** |
| Treatment (Control) |  |  |  |  |
| *Females* | -0.67 | 0.31 |  |  |
| *Males* | -0.12 | 0.28 |  |  |
| Treatment (None) |  |  |  |  |
| *Females* | -0.06 | 0.23 |  |  |
| *Males* | -0.16 | 0.18 |  |  |
| Foraging success x Mass | 0.036 | 0.06 | 0.57 | 0.57 |
| **Age x Mass** |  |  |  |  |
| *Females* | **-0.47** | **0.09** | **-5.08** | **< 0.0001** |
| *Males* | **-0.29** | **0.08** | **-3.79** | **0.0001** |
| Age x Sex (M) | 0.28 | 0.23 | 1.18 | 0.24 |
| Mass x Sex (M) | 0.22 | 0.17 | 1.26 | 0.21 |
| Group size x Sex (M) | 0.07 | 0.1 | 0.77 | 0.44 |
| Treatment (Control) x Sex (M) | 0.55 | 0.42 | 1.31 | 0.19 |
| Treatment (None) x Sex (M) | -0.1 | 0.28 | -0.36 | 0.72 |
| **Age x Treatment (Control)** |  |  |  |  |
| *Females* | **-0.62** | **0.21** | **-2.89** | **0.0039** |
| *Males* | -0.52 | 0.27 | -1.92 | 0.055 |
| Age x Treatment (None) |  |  |  |  |
| *Females* | 0.34 | 0.18 | 1.88 | 0.059 |
| *Males* | -0.006 | 0.16 | -0.03 | 0.97 |
| Age x Mass x Sex | 0.18 | 0.12 | 1.5 | 0.13 |
| Age x Treatment (Control) x Sex (M) | 0.09 | 0.34 | 0.28 | 0.78 |
| Age x Treatment (None) x Sex (M) | -0.34 | 0.24 | -1.45 | 0.14 |

Data used in these analyses were 182 observations of relative babysitting contributions to 28 litters produced in 9 groups across 3 years recorded from 105 subordinate meerkats.

**Figure 2.**
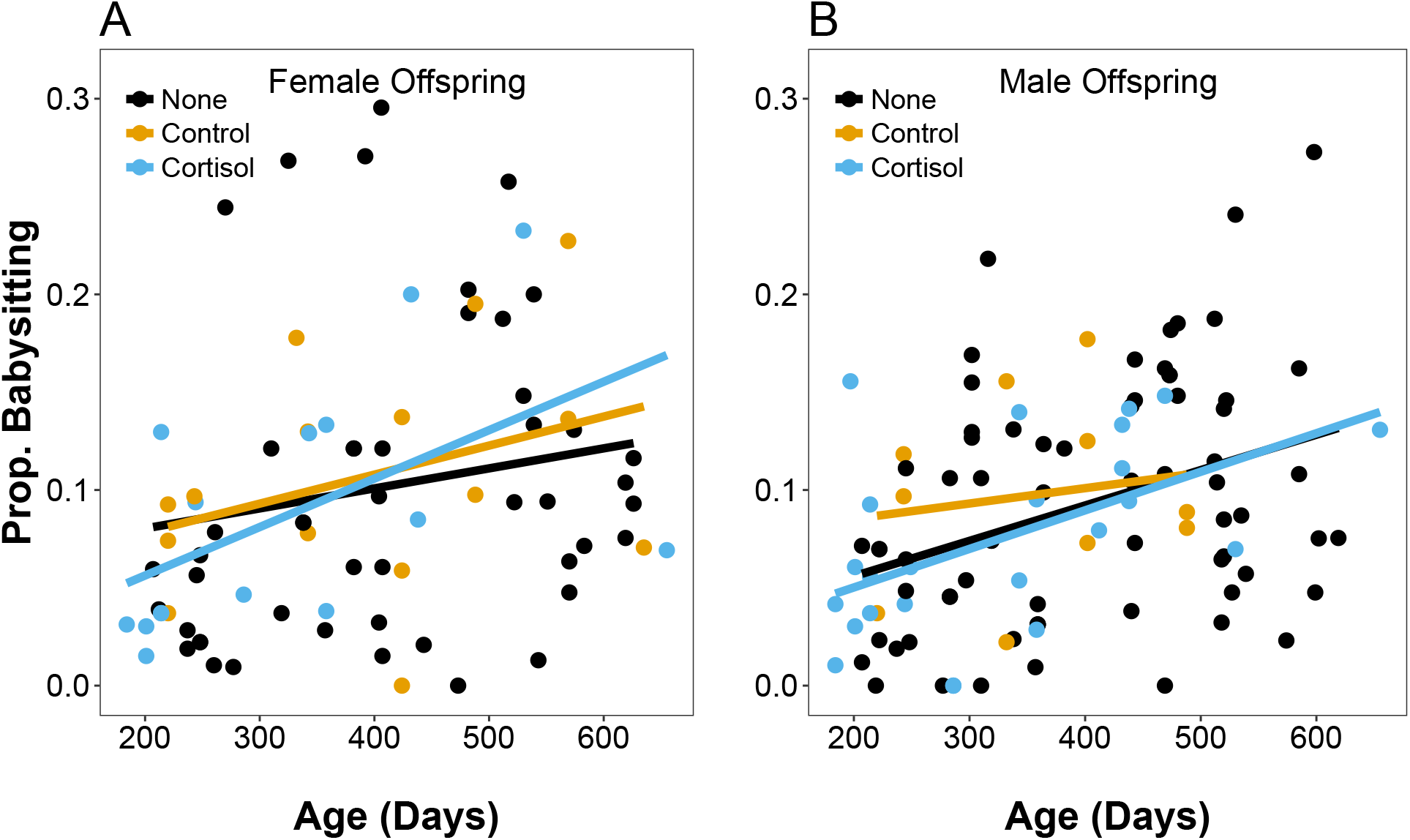
Babysitting contributions of (A) daughters and (B) sons from mothers treated with cortisol during pregnancy increased with age at a faster rate than those from control (females: *z* = −2.89, *P* = 0.0039; males: *z* = −1.92, *P* = 0.055), but not untreated (“None”) mothers (females: *z* = 1.88, *P* = 0.06; males: *z* = −0.03, *P* = 0.97, Table 4). Data are relative babysitting contributions from offspring from cortisol-treated (females: n = 15 estimates; males: n = 24), control (females: n = 15; males: n = 10), and untreated mothers (females: n = 49; males: n = 69). Raw data and regression lines are shown (full results in Table 4).

The effects of the maternal treatments on pup feeding depended upon the sex of the offspring, but not their age (Table 5). Daughters, but not sons from mothers treated with cortisol during pregnancy exhibited significantly more pup feeding contributions than those from control mothers (females: *z* = −3.12, *P* = 0.00018; males: *z* = −1.14, *P* = 0.25) or untreated mothers (females: *z* = −3.49, *P* = 0.0005, sons: *z* = −1.03, *P* = 0.3, Table 5, Fig. 3). Notably, the magnitude of effect size of maternal treatment for daughters was much larger than other variables known to impact babysitting contributions such as foraging success, age-related body mass, and group size (Table 5).

**Table 5.** Effect of dominant female treatments on relative pup feeding contributions. Data are from a generalized linear mixed-effects model where the response variable is the proportion of pup feeds exhibited by the subordinate meerkat relative to the total pup feeds the litter received. The model contained random intercept terms for individual (σ^2^ = 0.000), litter nested within group (σ^2^ = 0.2), year (σ^2^ = 0.08), and an observational level random intercept term to control for overdispersion (σ^2^ = 0.19). If fixed effects by themselves were involved in significant higher order interactions with other variables, only parameter estimates are shown.

| **Fixed Effect** | **b** | **SE** | **z** | **P-value** |
| --- | --- | --- | --- | --- |
| **Intercept** |  |  |  |  |
| *Females* | **-2.27** | **0.28** | **-8.2** | **<0.0001** |
| *Males* | **-2.66** | **0.25** | **-10.45** | **<0.0001** |
| **Observation time** | **0.59** | **0.06** | **9.41** | **<0.0001** |
| Litter size | -0.055 | 0.1 | -0.55 | 0.58 |
| Mixed litter (Y) | -0.03 | 0.29 | -0.1 | 0.92 |
| Sex (M) | -0.39 | 0.2 |  |  |
| Age |  |  |  |  |
| *Females* | -0.37 | 0.16 |  |  |
| *Males* | 0.29 | 0.14 |  |  |
| **Foraging success** | **0.14** | **0.06** | **2.12** | **0.033** |
| Mass |  |  |  |  |
| *Females* | 0.1 | 0.11 |  |  |
| *Males* | -0.37 | 0.11 |  |  |
| **Group size** |  |  |  |  |
| *Females* | **-0.48** | **0.13** | **-3.57** | **0.0003** |
| *Males* | **-0.32** | **0.13** | **-2.46** | **0.014** |
| **Treatment (Control)** |  |  |  |  |
| *Females* | **-0.73** | **0.23** | **-3.12** | **0.0018** |
| *Males* | -0.24 | 0.21 | -1.14 | 0.25 |
| **Treatment (None)** |  |  |  |  |
| *Females* | **-0.64** | **0.18** | **-3.49** | **0.0005** |
| *Males* | -0.14 | 0.13 | -1.03 | 0.3 |
| Foraging success x Mass | 0.06 | 0.06 | 0.97 | 0.33 |
| **Age x Mass** |  |  |  |  |
| *Females* | -0.04 | 0.07 | -0.62 | 0.54 |
| *Males* | **-0.19** | **0.07** | **-2.68** | **0.007** |
| **Age x Sex (M)** | **0.66** | **0.19** | **3.49** | **0.00048** |
| **Mass x Sex (M)** | **-0.47** | **0.13** | **-3.63** | **0.0003** |
| Group size x Sex (M) | 0.16 | 0.08 | 1.89 | 0.059 |
| Treatment (Control) x Sex (M) | 0.49 | 0.31 | 1.58 | 0.11 |
| **Treatment (None) x Sex (M)** | **0.5** | **0.22** | **2.23** | **0.023** |
| Age x Treatment (Control) |  |  |  |  |
| *Females* | -0.11 | 0.22 | -0.5 | 0.61 |
| *Males* | 0.03 | 0.29 | 0.11 | 0.91 |
| Age x Treatment (None) |  |  |  |  |
| *Females* | 0.17 | 0.16 | 1.03 | 0.3 |
| *Males* | -0.02 | 0.14 | -0.12 | 0.9 |
| Age x Mass x Sex | -0.14 | 0.09 | -1.51 | 0.13 |
| Age x Treatment (Control) x Sex (M) | 0.14 | 0.34 | 0.42 | 0.67 |
| Age x Treatment (None) x Sex (M) | -0.19 | 0.21 | -0.91 | 0.36 |

Data used in these analyses were 192 observations of relative pup feeding contributions to 26 litters produced in 7 groups across 3 years recorded from 101 subordinate meerkats.

**Figure 3.**
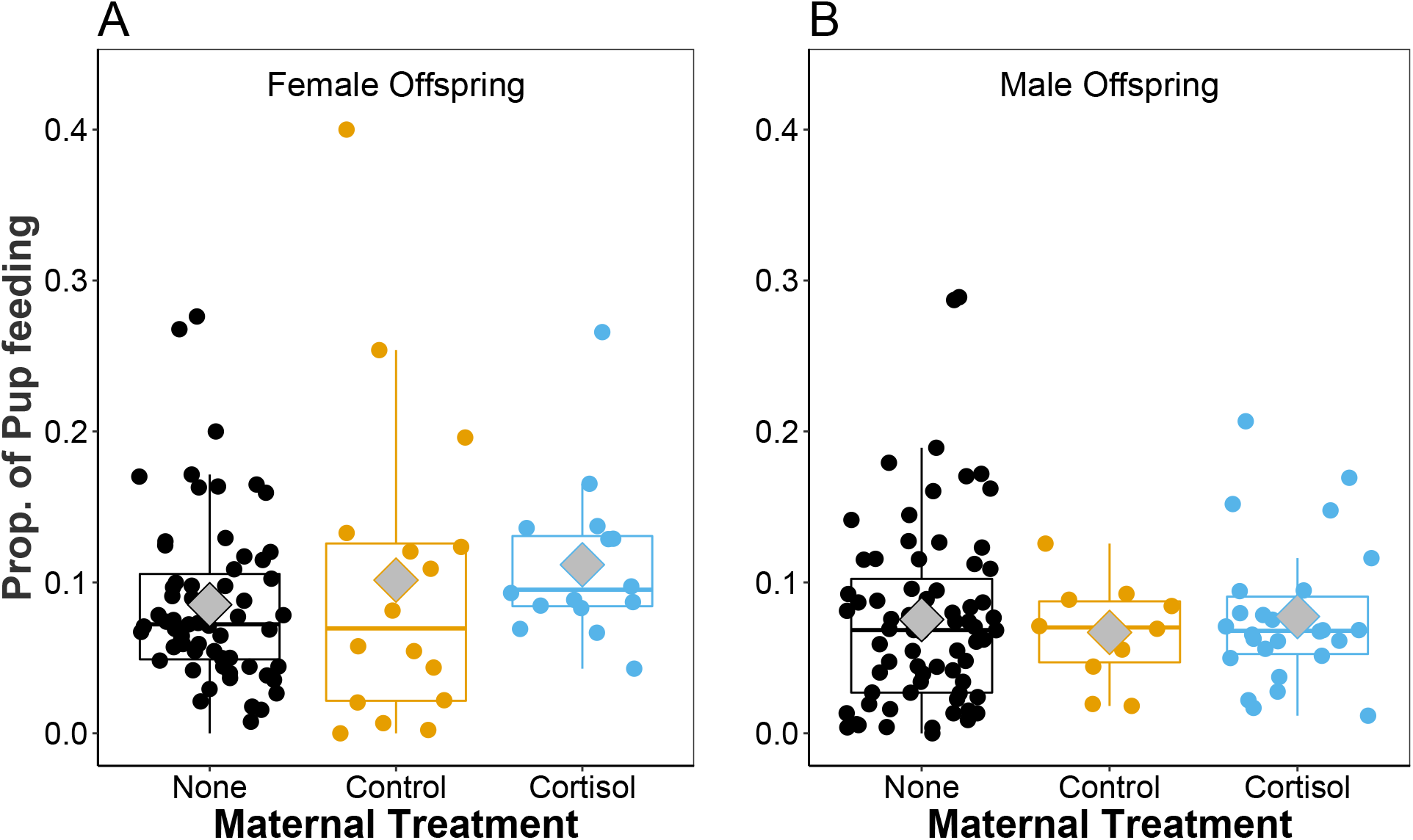
Pup feeding contributions of (A) daughters, but not (B) sons, from mothers treated with cortisol during pregnancy were significantly higher compared to from control (females: *z* = −3.09, *P* = 0.0004; males: *z* = −1.15, *P* = 0.25) or untreated (“None”) mothers (females: *z* = −3.47, *P* = 0.0005; males: *z* = −0.89, *P* = 0.37, Table 5). Data are relative pup feeding contributions from offspring from cortisol-treated (females: n = 16 estimates; males: n = 26), control (females: n = 16; males: n = 10), and untreated mothers (females: n = 64; males: n = 71). Raw data are shown (full results in Table 5). Boxplots show median (solid horizontal line), mean (grey diamonds), and first (25%) and third (75%) quartiles.

### Effects of treatments on offspring stress physiology

Daughters from mothers treated with cortisol during pregnancy had lower plasma cortisol concentrations (age x treatment, t = −1.76, P = 0.08, Table 6, Fig. 4A) and lower fGCM concentrations (age x treatment, t = −2.9, P = 0 .004, Table 7, Fig. 5A) as they became older compared to those from control mothers but these differences were only significant for fGCM concentrations. Sons from mothers treated with cortisol during pregnancy had significantly lower plasma cortisol concentrations as they became older compared to those from control mothers (age x treatment t = −2.68, P = 0.008, Table 6, Fig. 4B) but similar fGCM concentrations compared to those from control mothers as they became older (age x treatment, t = −0.1, P = 0.49, Table 7, Fig. 5B).

**Table 6.** Effect of dominant female treatments on plasma cortisol concentrations. Data are from a linear mixed-effects model where the response variable is plasma cortisol concentrations (ln transformed) of the subordinate meerkat. The model contained random intercept terms for individual nested within their birth litter (σ^2^ = 0.034), and capture group (σ^2^ = 0.000). If fixed effects by themselves were involved in significant higher order interactions with other variables, only parameter estimates are shown.

| Fixed Effect | b | SE | t | df | P-value |
| --- | --- | --- | --- | --- | --- |
| **Intercept** |  |  |  |  |  |
| *Females* | **3.18** | **0.29** | **11.31** | **128** | **<0.0001** |
| *Males* | **3.37** | **0.25** | **13.37** | **113** | **<0.0001** |
| **Sampling time** | **1.16** | **0.10** | **11.95** | **259** | **<0.0001** |
| **Sampling time<sup>2</sup>** | **-0.24** | **0.03** | **-7.24** | **269** | **<0.0001** |
| Time of day | -0.2 | 0.14 | -1.38 | 276 | 0.17 |
| Sample year (2015) | 0.41 | 0.24 | 1.72 | 228 | 0.09 |
| Sex (M) | 0.2 | 0.2 | 0.97 | 39 | 0.34 |
| Age |  |  |  |  |  |
| *Females* | -0.24 | 0.18 |  |  |  |
| *Males* | -0.01 | 0.14 |  |  |  |
| Foraging success |  |  |  |  |  |
| *Females* | 0.13 | 0.11 | -1.22 | 231 | 0.22 |
| *Males* | -0.15 | 0.1 | -1.42 | 269 | 0.16 |
| Group size | 0.07 | 0.11 | 0.62 | 242 | 0.54 |
| Group size<sup>2</sup> | 0.05 | 0.08 | 0.63 | 211 | 0.53 |
| Pups in group | -0.31 | 0.18 | -1.79 | 276 | 0.075 |
| **Group sex ratio** | **0.2** | **0.08** | **2.53** | **180** | **0.01** |
| Relatedness | -0.09 | 0.18 | -0.5 | 40 | 0.62 |
| Weather (PC1) | -0.08 | 0.1 | -0.75 | 276 | 0.45 |
| Treatment (Cortisol) |  |  |  |  |  |
| *Females* | -0.02 | 0.24 |  |  |  |
| *Males* | -0.32 | 0.19 |  |  |  |
| Sex (M) x Age | 0.22 | 0.22 | 1.04 | 272 | 0.3 |
| Sex (M) x Foraging success | -0.013 | 0.14 | -0.09 | 257 | 0.93 |
| Sex (M) x Treatment (Cortisol) | -0.3 | 0.3 | -1.01 | 35 | 0.32 |
| **Age x Treatment (Cortisol)** |  |  |  |  |  |
| *Females* | -0.42 | 0.24 | -1.76 | 276 | 0.08 |
| *Males* | **-0.5** | **0.19** | **-2.68** | **274** | **0.008** |
| Group size x Pups Present (Yes) | -0.05 | 0.15 | -0.36 | 216 | 0.72 |
| Group size<sup>2</sup> x Pups Present (Yes) | 0.07 | 0.12 | 0.6 | 263 | 0.55 |
| Group size x Weather (PC1) | -0.02 | 0.08 | -0.30 | 273 | 0.76 |
| Age x Sex (M) x Treatment (Cortisol) | -0.08 | 0.31 | -0.26 | 275 | 0.8 |

Data used in these analyses were 299 measures of plasma cortisol concentrations from 49 subordinate meerkats produced in 14 litters from 10 different groups. Reference levels (intercept) for “Sex” was female, “Pups in group” was Yes, and for Relatedness was “No parent was dominant”.

**Figure 4.**
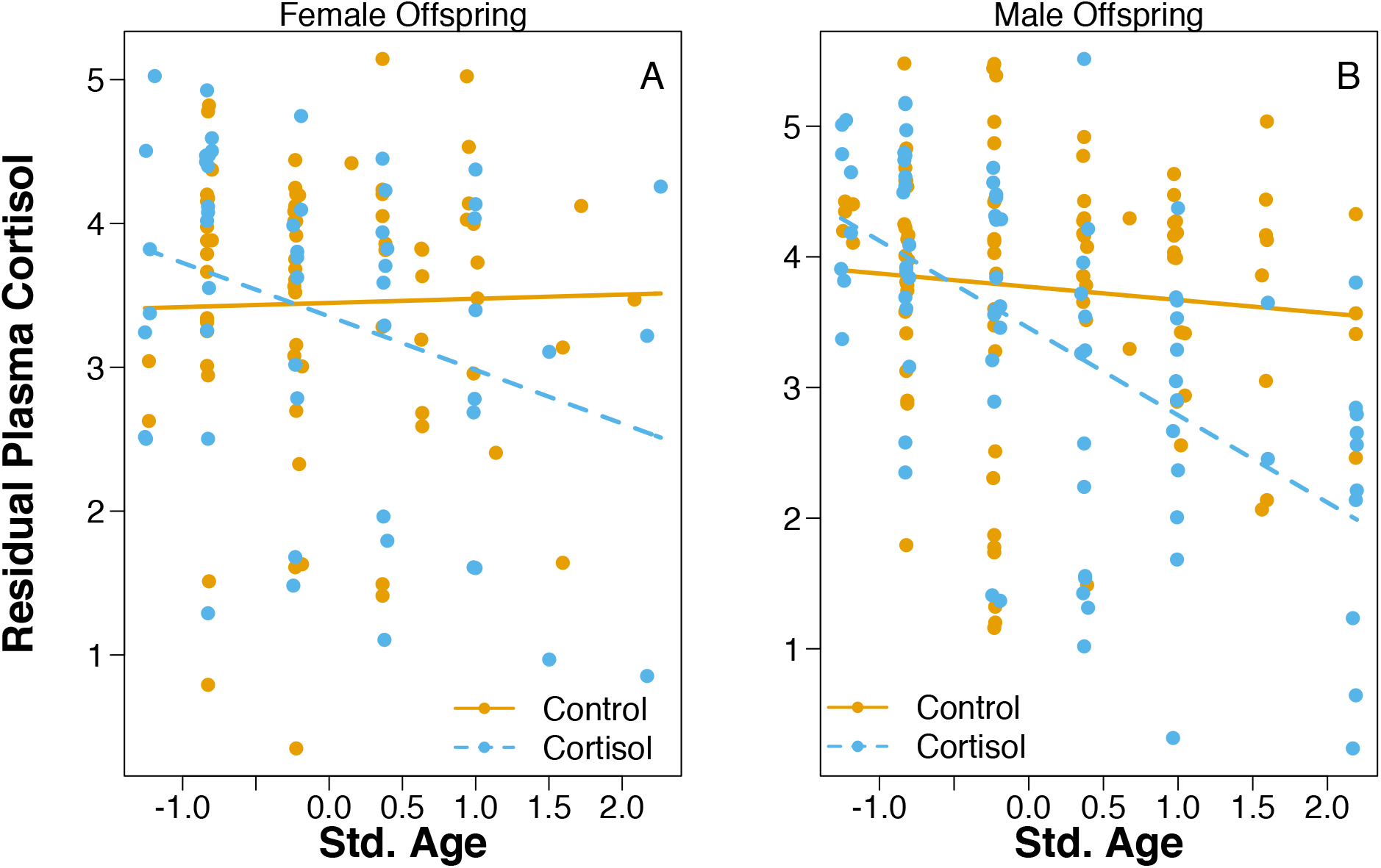
(A) Daughters and (B) sons from mothers treated with cortisol during pregnancy had lower plasma cortisol concentrations as they became older compared to those from control mothers, though the difference was only significant in males (daughters: age x treatment, t = −1.76, P = 0.08; sons: age x treatment t = −2.68, P = 0.008, Table 6). Data are residual plasma cortisol concentrations from offspring from cortisol-treated (females: n = 64 samples; males: n = 92) and control (females: n = 89; males: n = 104) mothers. Residuals from a linear mixed-effects model (Table 6) are shown on y-axis.

**Table 7.** Effect of dominant female treatments on faecal glucocorticoid metabolite (fGCM) concentrations. Data are from a linear mixed-effects model where the response variable fGCM concentrations (ln+1 transformed) of the subordinate meerkat. The model contained random intercept terms for individual nested within their birth litter (σ^2^ = 0.094) and collection group (σ^2^ = 0.000). If fixed effects by themselves were involved in significant higher order interactions with other variables, only parameter estimates are shown.

| **Fixed Effect** | **b** | **SE** | **t** | **df** | **P-value** |
| --- | --- | --- | --- | --- | --- |
| **Intercept** |  |  |  |  |  |
| *Females* | **5.4** | **0.25** | **21.36** | **50** | **<0.0001** |
| *Males* | **5.33** | **0.23** | **23.4** | **23** | **<0.0001** |
| **Time of day** | **-0.14** | **0.04** | **-3.29** | **516** | **0.001** |
| Sample year (2015) | 0.07 | 0.17 | 0.43 | 120 | 0.67 |
| Sex (M) | -0.06 | 0.2 | -0.31 | 22 | 0.76 |
| Age |  |  |  |  |  |
| *Females* | 0.48 | 0.16 |  |  |  |
| *Males* | 0.06 | 0.1 |  |  |  |
| Foraging success |  |  |  |  |  |
| *Females* | 0.16 | 0.13 | 1.29 | 355 | 0.2 |
| *Males* | 0.04 | 0.06 | 0.81 | 520 | 0.42 |
| **Group size** | **0.41** | **0.12** | **3.52** | **96** | **0.0007** |
| Group size<sup>2</sup> | -0.05 | 0.11 | -0.78 | 267 | 0.43 |
| Pups in group | -0.13 | 0.16 | -0.78 | 267 | 0.43 |
| **Group sex ratio** | **0.16** | **0.07** | **2.27** | **158** | **0.024** |
| Relatedness | 0.28 | 0.25 | 1.1 | 14 | 0.29 |
| Weather (PC1) | 0.09 | 0.07 | 1.4 | 203 | 0.16 |
| Treatment (Cortisol) |  |  |  |  |  |
| *Females* | -0.12 | 0.29 |  |  |  |
| *Males* | -0.03 | 0.24 |  |  |  |
| **Sex (M) x Age** | **-0.42** | **0.17** | **-2.47** | **221** | **0.014** |
| Sex (M) x Foraging success | -0.12 | 0.13 | -0.87 | 375 | 0.38 |
| Sex (M) x Treatment (Cortisol) | 0.1 | 0.33 | 0.29 | 18 | 0.77 |
| **Age x Treatment (Cortisol)** |  |  |  |  |  |
| *Females* | **-0.63** | **0.22** | **-2.9** | **447** | **0.004** |
| *Males* | -0.1 | 0.14 | -0.69 | 491 | 0.49 |
| Group size x Pups Present (Yes) | -0.17 | 0.12 | -1.46 | 216 | 0.14 |
| Group size<sup>2</sup> x Pups Present (Yes) | 0.11 | 0.12 | 456 | 0.97 | 0.33 |
| Group size x Weather (PC1) | 0.04 | 0.05 | 0.69 | 129 | 0.49 |
| **Age x Sex (M) x Treatment (Cortisol)** | **0.53** | **0.25** | **2.15** | **413** | **0.032** |

Data used in these analyses were 542 faecal samples (n= 355 from controls, n = 187 from cortisol treated litters) from 34 subordinate meerkats (control: n = 12 females, n = 11 males; cortisol-treated: n = 5 females, n = 6 males) produced in 10 litters from 7 different groups. Reference levels (intercept) for “Sex” was female, “Pups in group” was Yes, and for Relatedness was “No parent was dominant”.

**Figure 5.**
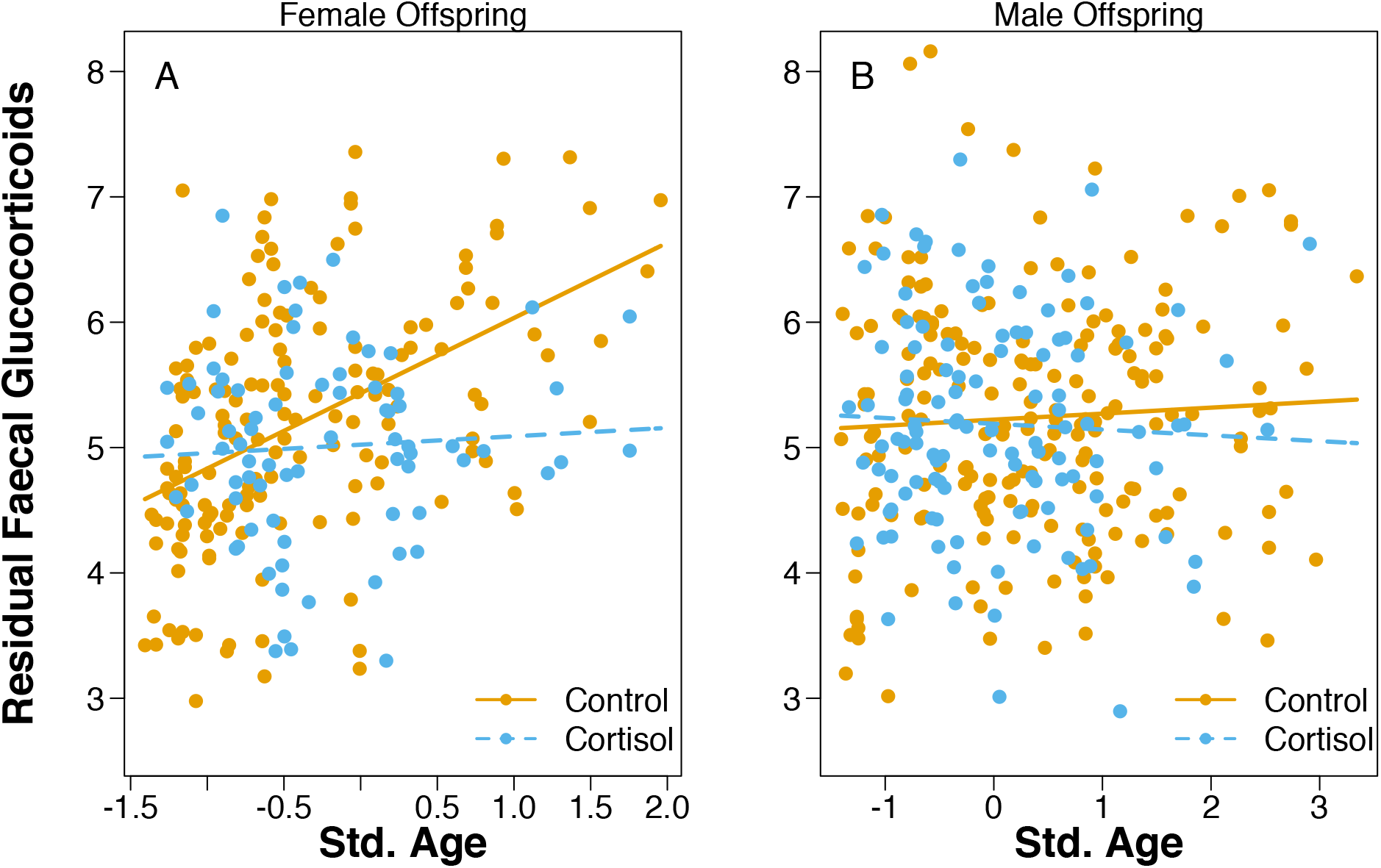
Faecal glucocorticoid metabolite (fGCM) concentrations in (A) daughters but not (B) sons from mothers treated with cortisol during pregnancy were significantly lower than those from control mothers as they became older (daughters: age x treatment, t = −2.9, P = 0.004; sons: age x treatment, t = −0.1, P = 0.49, Table 7. Data are residual fGCM concentrations from offspring from cortisol-treated (females: n = 79 samples; males: n = 118) and control (females: n = 154; males: n = 201) mothers. Residuals from a linear mixed-effects model (Table 7) are shown on y-axis.

## Discussion

We found some support for our hypothesis that elevated maternal GCs would reduce the potential for offspring to have direct reproductive opportunities and would therefore shift them towards exhibiting more cooperative behaviour that could increase their indirect fitness. Daughters, but not sons, from mothers treated with cortisol during pregnancy grew more slowly early in life and exhibited more babysitting and pup feeding behaviour as they became older compared to controls. Other than offspring survival (Table 2), we were unable to quantify the direct and indirect fitness of offspring from control or cortisol-treated mothers, but early life growth or body mass (which we measured here) is closely linked to direct fitness opportunities in daughters (27-32). Previous studies in meerkats show that female, but not male, offspring that grow faster from 1-3 months are more likely to acquire the dominant breeding position (31), perhaps because offspring that grow faster in their first 3 months of life are heavier later in life (32, 57, 58), and heavier females are more likely to acquire a vacant dominant breeding position (30, 32). As such, daughters, but not sons, from mothers treated with cortisol levels during pregnancy should have reduced future direct fitness opportunities and therefore increase their investment in behaviours that elevate their indirect fitness. Our results are consistent with studies in other taxa that suggest that individuals adjust their contributions to cooperative behaviour according to their future reproductive potential. For example, in cooperatively breeding birds, when the chances of direct reproduction are elevated, subordinates often stop helping at the nest (59). Studies of social wasps show that individuals whose probability of acquiring the dominant breeding position was experimentally increased exhibited significantly less helping behaviour (60, 61). Finally, in cooperatively breeding fish, subordinates will reduce their helping investment immediately prior to dispersal from their natal group where they attempt to reproduce on their own rather than stay in their natal group and queue for dominance (62).

Our results show that increases in maternal GCs can increase the cooperative behaviour of daughters, which should lead to substantial direct fitness benefits to mothers. Daughters from mothers treated with cortisol during pregnancy exhibited more alloparental care compared to controls, such that subsequent offspring produced in groups with offspring from cortisol-treated mothers should have received more alloparental care. Because offspring that receive more alloparental care grow faster early in life or are larger later in life (32, 57), the presence of offspring from cortisol-treated mothers should increase the direct fitness of dominant breeders and the indirect fitness of the offspring from cortisol-treated mothers. Taken together, our results suggest that this GC-mediated maternal effect reduced the direct fitness opportunities of daughters by reducing their early life growth, but they compensated by increasing their investment in indirect fitness opportunities (helping to rear non-descendent offspring).

This is in line with theoretical predictions that parental manipulation of the cooperative behaviour of offspring can evolve if the costs of resisting the parental effect are high and inclusive fitness benefits of helping rear subsequent offspring are increased (18), as is the case in cooperative breeders.

Control females that were fed during pregnancy produced daughters that grew faster during early development (1-3 months) compared to daughters from cortisol-treated or untreated mothers. Although mothers that were treated with cortisol during pregnancy received the same amount of supplemental food as controls, daughters and sons from mothers fed cortisol during pregnancy did not differ in early life growth compared to those from untreated mothers. This indicates that the additional food provided to dominant females during pregnancy had the potential to increase growth, but the added cortisol prevented those gains in body mass. This has implications for understanding the fitness consequences of maternal stress on offspring growth trajectories (15, 63) because our results show that elevated circulating GC levels in pregnant females in the absence of energetic constraints induced reductions in the early life growth of offspring. This supports the hypothesis that maternal GC levels during offspring development act as a cue that induces plasticity in offspring growth rather than simply mediating the effects of energetic constraints. Alternatively, elevated maternal GCs could alter patterns of maternal investment in offspring. Identifying whether offspring or mothers are driving these effects is a major challenge in studies of maternal stress effects in wild animals.

The reductions in the activity of the neuroendocrine stress axis of daughters may have potentiated the increased alloparental care behaviour that we observed. Compared to daughters from control mothers, daughters from mothers treated with cortisol during pregnancy exhibited more babysitting as they became older, more overall pup feeding, and they also had lower plasma cortisol and fGCM concentrations. Males from mothers treated with cortisol during pregnancy had significantly lower plasma cortisol concentrations, but not fGCM concentrations as they got older and also tended to exhibit more babysitting as they aged. The activity of the neuroendocrine stress axis is closely linked to an array of social behaviours (64) and our recent work shows that elevated activity of the neuroendocrine stress axis reduces babysitting in both females and males and decreases pup feeding in females (39). Together, this supports the hypothesis that the mechanism by which early life stress increases the cooperative behaviour of daughters is by dampening the activity of their neuroendocrine stress axis.

Our results show that the effects of maternal GCs on offspring growth, physiology, and behaviour were greater in daughters than in sons, which adds to biomedical (65-66) and ecological (67-69) studies that highlight how early life conditions or maternal GC levels can have sex-specific consequences for offspring. In meerkats, there may be added benefits for the dominant female for altering the cooperative behaviour of daughters compared to sons; daughters exhibit more cooperative behaviour than sons (36) and are more responsive to the begging calls of subsequent offspring that they provision with food (70). More broadly, sex-differences in natal dispersal may cause these differential responses to parental effects. In meerkats, subordinate males voluntarily disperse from their natal group to look for receptive females but can return to their natal group whereas subordinate females rarely voluntarily disperse from their natal group (71). In our case and in others (63), the more philopatric sex (females) is more sensitive to early life conditions, which may be due to differential costs of parental modification between the philopatric and dispersing sex. If parental effects have long-term consequences on offspring characteristics, as we show here, there may be an increased degree of mismatch between the phenotype of the dispersing sex and the postnatal environment where individuals eventually settle. If this mismatch has fitness costs, this should select for individuals from the dispersing sex to be less responsive to cues from the parental phenotype or environment.

Our results provide some support for the hypothesis that parents may alter the cooperative tendencies of their offspring by manipulating the characteristics of their offspring (9-10), though we note that it is uncertain if the transfer of maternal GCs to offspring was passive or active. Explanations regarding the evolutionary origins of cooperative behaviour involve nepotism or kin selection (72), mutualisms or reciprocity (73), but few studies have tested the “parental manipulation” hypothesis proposed by Alexander (9). Some studies show that alleles that increase maternal fitness at the expense of the direct fitness of offspring can evolve (19) and that cooperative breeders may bias investment towards offspring that exhibit more cooperative behaviour (74). Our study supports the hypothesis that environmental stressors may induce a parental effect that can modify the cooperative tendencies of their offspring.

Finally, our results have two implications for theoretical models examining the evolution of parental effects. First, given the sex-specificity of parental effects, our results challenge the conclusions of models examining the evolution of parental effects that assume that all offspring are equally sensitive to the parental effect (16), or those that assume that the benefits of exhibiting the phenotype resulting from the parental effect are equal for all offspring (18). Second, selfish parental effects are thought to be relatively rare (8, 13) and theory (16-17) and empirical studies showing sex-specific responses to early life stress (65-66) indicate that offspring can become resistant to such selfish parental effects. However, some models indicate that the evolution of selfish parental effects may be dependent upon the social environment (24), especially if the selfish parental effect influences the expression of alloparental care behaviour of offspring and therefore increases the indirect fitness of offspring. Our results provide an example whereby a GC-mediated maternal effect should decrease the direct fitness of daughters (by reducing their early life growth), but should increase the direct fitness of mothers and indirect fitness of daughters by elevating their cooperative behaviour.

### Ethics

All protocols used in our experiments were approved by the Animal Ethics Committee at the University of Pretoria (Pretoria, South Africa: #EC031-13, #EC047-16) and the Northern Cape Department of Environment and Nature Conservation Research (South Africa: FAUNA 050/2013, FAUNA 192/2014, FAUNA 1020/2016).

## Acknowledgements

We thank M. Manser for her contributions to maintain the long-term Kalahari Meerkat Project (KMP) and her contributions to this specific project. Thanks to P. Roth, S. Bischoff-Mattson, L. Howell, J. Samson, N. Thavarajah, and numerous KMP volunteers who contributed to data collection. Thanks to M. Manser and S. Guindre-Parker for helpful comments. Thanks to the Northern Cape Conservation Authority for permission to conduct research and the Kotze family for allowing us to study meerkats on their property.

## Authors’ Contributions

B.D. and T.H.C-B planned the study, B.D. designed the experiments, B.D., C. Dubuc, D.L.C, T.H.C-B, D.G., I.BG., N.B., M.H., A.G., and C. Duncan coordinated and collected data, B.D. conducted analyses and produced figures, B.D. and T.H.C-B authored manuscript with contributions from all authors.

## Date, code and materials

All data will be archived on Data Dryad. All code for statistical analyses is available from first author.

